# Postnatal functional inactivation of the ventral subiculum enhances dopaminergic responses in the core part of the nucleus accumbens following ketamine injection in adult rats

**DOI:** 10.1101/859322

**Authors:** Hana Saoud, Duco De Beus, Séverine Eybrard, Alain Louilot

## Abstract

For almost two decades schizophrenia has been considered to be a functional disconnection disorder. This functional disconnectivity between several brain regions could have a neurodevelopmental origin. Various approaches suggest the ventral subiculum (SUB) is a particular target region for neurodevelopemental disturbances in schizophrenia. It is also commonly acknowledged that there is a striatal dopaminergic (DA) dysregulation in schizophrenia which may depend on a subiculo-striatal disconnection involving glutamatergic NMDA receptors.

The present study was designed to investigate, in adult rats, the effects of the non-competitive NMDA receptor antagonist ketamine on DA responses in the ventral striatum, or, more specifically, the *core* part of the nucleus accumbens (Nacc), following postnatal functional inactivation of the SUB. Functional inactivation of the left SUB was carried out by local tetrodotoxin (TTX) microinjection at postnatal day 8 (PND8), i.e. at a critical point in the neurodevelopmental period. DA variations were recorded using *in vivo* voltammetry in freely moving adult rats (11 weeks). Locomotor activity was recorded simultaneously with the extracellular levels of DA in the *core* part of the Nacc. Data obtained during the present study showed that after administration of ketamine, the two indexes were higher in TTX animals than PBS animals, the suggestion being that animals microinjected with TTX in the left SUB at PND8 present greater reactivity to ketamine than animals microinjected with PBS. These findings could provide new information regarding the involvement of NMDA glutamatergic receptors in the *core* part of the Nacc in the pathophysiology of schizophrenia.

## 1. Introduction

Schizophrenia is a severe psychiatric illness, affecting about 0.7-0.8 % of the world population, commencing in young adulthood and evolving towards a chronic state (Saha et al., 2005). For about two decades now schizophrenia has been considered to be a functional disconnection disorder (Friston and Frith, 1995; Fornito and Bullmore, 2015; Cao et al., 2016). This functional disconnectivity between distributed regions of the brain might have a neurodevelopmental origin (Weinberger, 1995; Inta et al., 2011; Piper et al., 2012; Fornito and Bullmore, 2015; Owen et al., 2016). One of these regions, the parahippocampal structure, subiculum, would appear to be a particular target for neurodevelopemental disturbances in schizophrenia, as established by post-mortem data showing molecular, cellular, and morphological anomalies characteristic of developmental impairments (Arnold and Rioux, 2001). Moreover, recent neuroimaging studies in at-risk subjects have shown that with the onset and progression of the schizophrenia psychosis marked changes in terms of metabolism and structure are observed in the subiculum, in the left brain hemisphere (Schoebel et al., 2013).

Central to the pathophysiology of schizophrenia, and consistent with the common blockading action of antipsychotics on dopaminergic D2 type receptors, the hypothesis of a striatal DAergic dysfunction in schizophrenia was proposed several decades ago (Weinstein et al., 2017). However, the nature and/or origin of the DA dysfunction in schizophrenia has still not been fully elucidated. Moreover, recent brain imaging studies suggest that DA dysfunctioning does not follow an even distribution in the striatum. More precisely, the suggestion is that there is elevated presynaptic DAergic functioning (characterized by increased DA synthesis and enhanced phasic DA release) in the associative striatum but not, surprisingly, in the limbic striatum (ventral striatum) comprising, the Nacc as a whole (Weinstein et al., 2017). However, some differential changes involving the various sub-regions of the Nacc cannot be ruled out at this level insofar as since the 1990s the Nacc has been subdivided into two main sub-regions, the *core* and *shell* parts, which are anatomo-functionally distinct (Heimer et al., 1997). The proposal according to which subtle changes occur in Nacc sub-territories in schizophrenia is supported by recent post-mortem observations. Data suggest that in patients with schizophrenia the control of DA release by glutamatergic afferents might be more disrupted in the *core* part of the Nacc than in the *shell* part (McCollum et al., 2015). More specifically, post-mortem data obtained by McCollum et al. (2015) suggest that in patients with schizophrenia there is an excessive glutamatergic-type input in the Nacc, but only in the *core* sub-region. It is interesting in this connection to note that an increase in glutamate release has been described in the Nacc after phencyclidine or ketamine administrations (Adams and Moghaddam, 1998; Moghaddam and Adams, 1998), and that it has also been reported that these non-competitive NMDA antagonists induce typical symptoms of schizophrenia in healthy controls and increase such symptoms in patients with schizophrenia (Lahti et al., 2001; see Coyle et al, 2012).

Within the framework of animal modelling of the pathophysiology of schizophrenia in a heuristic perspective, and taking into account the afore-mentioned elements, the present study was designed to investigate the consequences of neonatal inactivation of the ventral subiculum (SUB) in adult rats in terms of behavioural (locomotor) responses and DAergic responses to ketamine challenge in the left *core* part of the Nacc. To be more precise, the well-known specific blocker of the pore of sodium channels, tetrodotoxin (TTX) (Moczydlowski, 2013), was microinjected in the SUB at post-natal day 8. As discussed previously (Meyer and Louilot, 2014), when locally administered during the postnatal developmental period TTX has been reported to affect myelin formation, the transformation of filiform spines into mature dendrite spines, and to alter normal axon branching and the final arrangement of synaptic connections. In rats, postnatal day 8 corresponds to the second trimester of pregnancy in humans, a period of great vulnerability for developing schizophrenia. Moreover, disruptions of myelin formation and dendritic spines have been found in patients with schizophrenia (see Meyer and Louilot, 2014; Tagliabue et al., 2017 for discussion). In other respects, the behavioural index chosen in the present study, namely locomotor activity, is generally proposed to have good translational relevance for psychotic symptoms in humans, given the increase in locomotor activity observed in rodents and the appearance of positive symptoms in humans after administration of NMDA antagonists or other psychotomimetic drugs (Winship et al., 2019). Changes in locomotor activity and DA levels in the *core* subregion of the Nacc were monitored in parallel, in freely moving rats, using *in vivo* voltammetry in adult animals.

## 2. Materials and methods

### 2.1. Animals and study design

All the experiments reported herein comply with the ARRIVE guidelines and were performed in accordance with Directive 2010/63/EU on the protection of animals used for scientific purposes. They were authorized by the Strasbourg Regional Ethics Committee on Animal Experimentation (CREMEAS-CEEA35) and the French Ministry of National Education, Superior Teaching, and Research (authorization APAFIS# 1553-2015082613521475 v3). Every effort was made to minimize the number of animals used and their suffering.

All the experiments were carried out on male Sprague-Dawley rats born to mothers obtained from Janvier-Labs (Le Genest St Isle, 53940, France). Throughout the experiment the animals were housed at +22 ± 2 °C, with food and water available *ad libitum*. Gestant mothers were housed individually in plexiglas cages before parturition. Neonates and mothers were maintained on a 12h:12h light-dark cycle (lights on at 7:00 am).

The design of the study is similar to that reported in previous articles (Meyer and Louilot, 2011; 2012; Tagliabue et al., 2017). Neonates’ day of birth was considered to be as postnatal day 0 (PND0). At postnatal day 8 (PND8), neonates randomly received PBS (controls) or TTX (experimental animals) microinjected in the ventral subiculum (SUB). From postnatal day 56 (PND56), male rats were accustomed to an inverted light-dark cycle (lights off from 11:00 am to 11:00 pm). At postnatal day 70 (PND70), a specially designed microsystem enabling the simultaneous measurement of DA and behaviour was surgically implanted in adult animals. After about one week of post-surgical recovery, implanted animals were subjected to the pharmacological experiment during the dark period of the inverted light-dark cycle.

### 2.2. Postnatal functional blockade of the SUB

Postnatal TTX functional blockade of the left SUB was performed at PND8. Surgery in neonates was conducted under inhalational anaesthesia, using the volatile anaesthetic, isoflurane (Isovet®, CSP, Cournon-d’Auvergne, France), as previously described (Meyer and Louilot, 2011; 2012; Usun et al., 2013). Neonatal microinjection of one of the two solutions, PBS or TTX (100μM), was then carried out, at random, in all the pups of one litter, by means of a stainless steel cannula (30 gauge, 12.5mm length, Small Parts, Miami, USA) implanted in the SUB with the following coordinates: 0.6 mm anterior to the interaural line (AP); 4.15 mm lateral to the midline (L); and 4.85 mm below the cortical surface (H). The volume of the microinjected solution was 0.3 μl, and the rate 2 min 15 s. The procedure has already been presented in detail in past papers (Meyer and Louilot, 2011; 2012; Usun et al., 2013). Thus, as previously discussed at length, it is unlikely under our conditions, which correspond to about 10 ng microinjected TTX (100μM × 0.3μl), that the radius of the efficient spread of TTX microinjected in the SUB would be greater than 0.67 mm, with TTX blockade action lasting 4h-48h.

### 2.3. Adult surgical implantation of the specially devised microsystem

A specially designed microsystem (Unimécanique-M2E, France; Louilot et al., 1987a), enabling behavioural and DAergic responses to be recorded simultaneously, was stereotaxically implanted at PND70 in the grown male rats, weight ∼400 ± 25 g, microinfused in the SUB at PND8 with either PBS or TTX. Stereotaxical surgery was performed using a stereotaxic frame (Unimécanique-M2E, France) following general anaesthesia with chloral hydrate (400 mg/kg; 8 ml/kg ip) (Field et al., 1993) and sc injection of a local anaesthetic (Xylovet®, Ceva, Libourne, France) in the animal’s head to ensure as full analgesia as possible throughout the implantation. The coordinates used for the implantation were: 10.6 mm (AP); 1.6 mm (L); and 7.1 mm (H). Animals were allowed a post-surgery recovery period of at least a week.

### 2.4. Behavioural analysis

The behavioural study, involving 73 animals in total, aimed to investigate the changes in locomotor activity following ketamine administration. Changes in locomotor activity and DA variations in the shell part of the Nacc were measured in parallel, before and after the sc injection of saline or ketamine at 0.5 ml/kg. Control animals were injected with saline (NaCl 0.9%), whereas the experimental animals were administered with ketamine (Imalgene®, Merial, Lyon, France) at 5 mg/kg,10 mg/kg and 20 mg/kg. Once the adult animals had recovered from the surgical operation, the pharmacological experiment could take place. The carbon fibre microelectrode was first positioned in the *core* part of the Nacc using the specially devised, implanted microsystem. Then the animal was placed in the experimental cage (27 cm long × 24 cm wide × 44 cm high). After about one hour of habituation to the cage, the animal randomly received the sc injection of either NaCl 0.9% or one of the 3 doses of ketamine and was kept in the experimental cage for a further 90 min. A small infrared camera placed in the top of the cage and connected up to a video system (monitor+ recorder) recorded the animal’s behaviour. The floor of the cage was divided into four virtual parts, all with the same surface area. To quantify locomotor activity, the number of times the animal moved from one quadrant to another (by engaging at least the head), as visually observed, was counted for 10-mins periods. Locomotor variations were expressed as mean±SEM.

### 2.5. Variations in the dopaminergic signal

The electrochemical *in vivo* detection of DA in the *core* part of the Nacc was achieved as previously described (Meyer and Louilot, 2011; 2012; Tagliabue et al., 2017). To be more precise, the voltammetric method used was differential normal pulse voltammetry (DNPV) combined with carbon fibre microelectrodes subjected to electrochemical pre-treatment and numerical mathematical analysis of the DNPV signal (Gonzalez-Mora et al., 1991). The mean value of the last 10 peaks of DA recorded during the control period (∼1h) before sc injection of NaCl 0.9% or one of the 3 doses of ketamine was calculated for each rat and taken as the 100% value. DA variations recorded every min were thus expressed as percentages (mean ± S.E.M.).

### 2.6. Statistics

Behavioural and voltammetric results were statistically analyzed using a multifactorial analysis of variance (ANOVA), with repeated measures on the time factor. Only between-subject ANOVAs are presented unless otherwise indicated. Between-subject variables were neonatal microinjection factor, with two levels (PBS or TTX), and ketamine dose factor, with four levels (NaCl 0.9% sc, ketamine 5mg/kg sc, ketamine 10 mg/kg sc, or ketamine 20 mg/kg sc). The dependent variables, for the behavioural study were the number of crossings per 10min-period, and for the voltammetric study the DA changes in the *core* part of the Nacc. Post-hoc contrast analyses of the ANOVA were carried out for the behavioural study to test specific hypotheses (Rosenthal et al., 2000). P<0.05 was the statistical significance level for all the analyses.

### 2.7. Histology

For the sake of histological verification of the location of the microinjection site in the SUB and the voltammetric recording site in the *core* part of the Nacc, animals were euthanized with a lethal injection of Dolethal® (5 ml/kg i.p.), 24-48h after the experiment. Their brains were removed from the skull and kept at +4°C in a 8% paraformaldehyde + 30% sucrose solution. The site of the neonatal microinjection in the SUB was located with the help of the Evans Blue vital dye previously added to the PBS and TTX solutions, and Neutral Red coloration of brain sections. The location of the recording site in the *core* was identified by electrocoagulation at the end of the pharmacological experiment, as previously described in detail (Meyer et al., 2009), and by the Thionin coloration of brain sections. The Paxinos and Watson atlas (2009) was used as a reference to identify the different brain structures.

## 3. Results

### 3.1. Histology (Figure 1)

Macroscopic qualitative examinations of the brain sections at the level of the SUB or *core* part of the Nacc showed no evidences of gliosis, or anatomical alterations, in animals microinjected in the SUB at PND8 with either PBS (Figure 1A) or TTX (Figure 1B). Only animals displaying microinjection sites clearly located in the left SUB (Figure 1, bottom) were taken into account for the locomotor activity and voltammetric analyses. Rats with a recording site outside the left *core* part of the Nacc (Figure 1, top) were only considered for the behavioural analysis, not for the voltammetric analysis.

**Figure 1.**
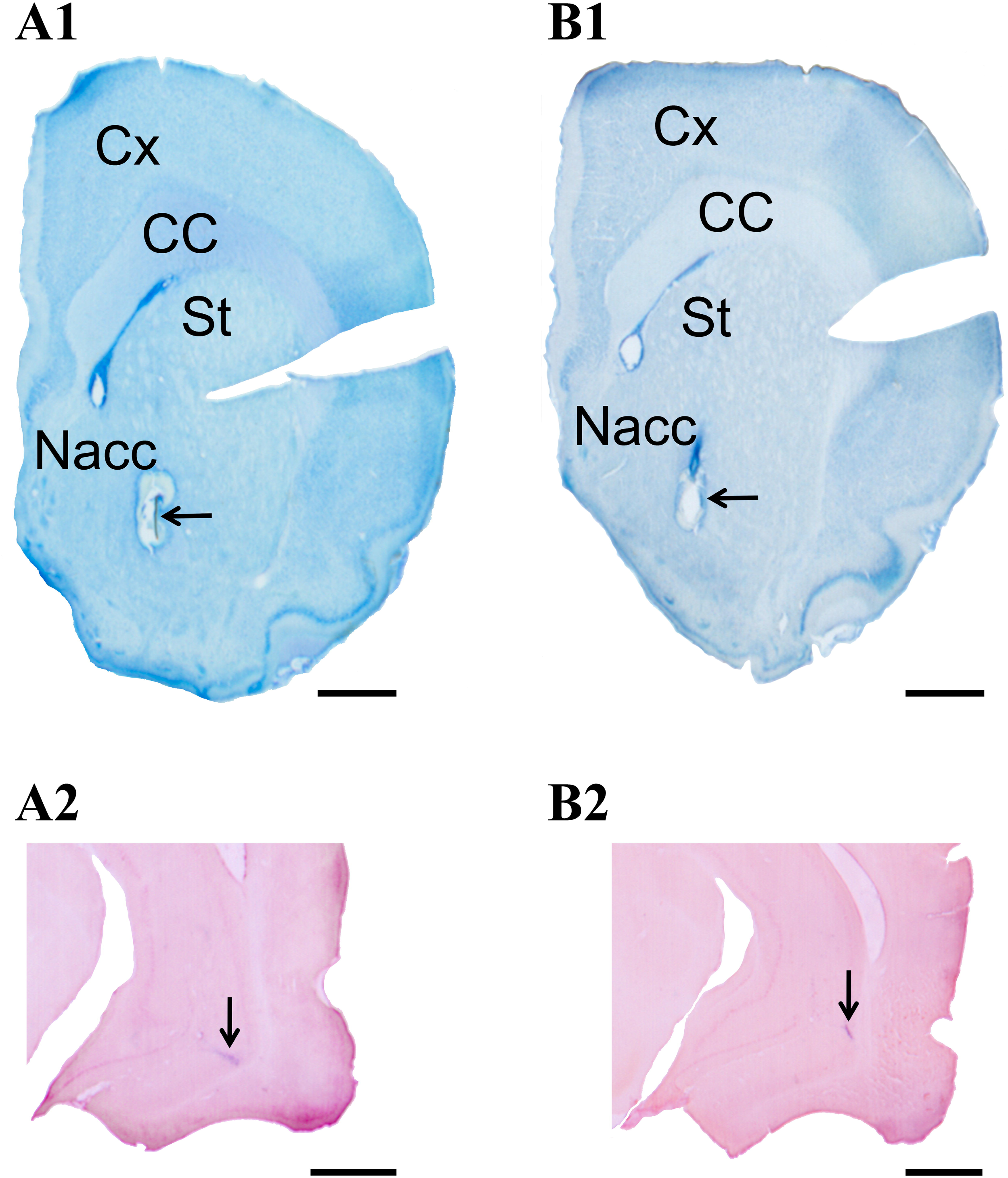
Brain sections in adult rats illustrating typical voltammetric recording sites in the left *core* subregion of the nucleus accumbens (A1, B1) and typical microinjection sites in the left ventral subiculum (A2, B2) of PBS (left microphotographs) or tetrodotoxin (TTX) (right microphotographs). The recording sites in the *core* subregion (A1, B1; arrows) are located by electrocoagulation carried out at the end of the experiment and Thionin staining. The neonatal microinjection sites (arrows) of PBS (A2) or TTX (B2) are identified by Evans Blue added to the PBS and TTX solutions microinjected in the ventral subiculum at postnatal day 8 and by Neutral Red staining of adult brain sections. Scale bar = 1 mm. Cx, cortex; CC, corpus callosum; Nacc, nucleus accumbens; St, striatum

### 3.2. Locomotor activity following administration of ketamine (Figure 2)

In total 73 animals were used to investigate ketamine-induced locomotor activity. Spontaneous locomotor activity (before the injection of saline or ketamine) in the different groups was not found to be statistically different. As regards the 10 min preceding the sc injection, the results provided by the ANOVA were F[1,65] = 0.67; ns for the postnatal microinjection, and F[3,65] = 0.54; ns for the dose x postnatal microinjection.

**Figure 2.**
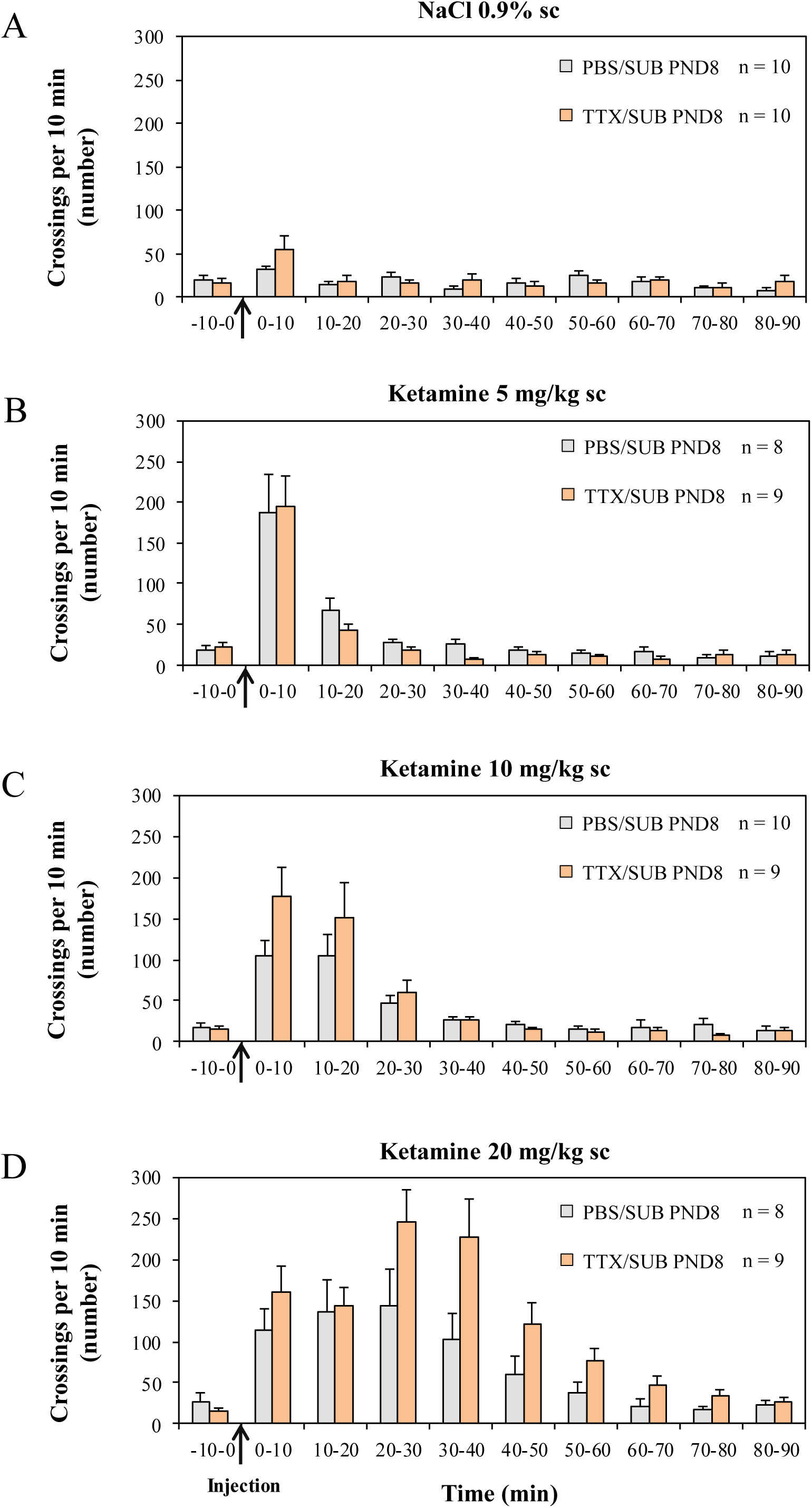
Ketamine-induced locomotor responses in adult rats microinjected with PBS (grey columns) or TTX (orange columns) at postnatal day 8 (PND8) in the ventral subiculum (SUB). Animals received a subcutaneous (sc) injection (arrow) of NaCl 0.9% (A), ketamine 5 mg/kg (B), ketamine 10 mg/kg (C), or ketamine 20 mg/kg (D). Locomotor activity is expressed as means ± SEM of the number of crossings per 10 min period. n is the number of animals in each group. Results are statistically analyzed using factorial ANOVA.

Statistical analysis showed that locomotor activity was dose-dependent and microinjection-dependent during the 90 min following saline and ketamine injections. Thus, the general ANOVA carried out for the 90 min post-injection displayed a significant dose-effect (NaCl 0.9%, ketamine 5 mg/kg, ketamine 10 mg/kg or ketamine 20 mg/kg) (F[3,65] = 24.54 p<0.000001), a significant postnatal microinjection-effect (PBS/TTX) (F[1,65] = 4.57 p<0.05) and a significant dose × microinjection interaction (F[3,65] = 2.91 p<0.05). Contrast analysis of ANOVA was performed for the 90 min following the injection to test the hypothesis that locomotor activity is statistically dependent on the postnatal microinjection (PBS/TTX) for the 2 higher doses of ketamine (10 mg/kg and 20 mg/kg). A significant microinjection effect (PBS/TTX) was found for the highest ketamine dose, 20 mg/kg s.c. (F[1,65] = 11.90 p<0.001), but not the 10 mg/kg s.c. ketamine dose (F[1,65] = 0.81 ns).

The effects of ketamine on the time course of locomotor activity for the 90 min post-injection period as established by within-subjects analysis were as follows: a significant effect of time (F [8,520] = 47.50 p < 0.00001) and a significant time × dose interaction (F [24,520] = 11.46 p < 0.00001). By contrast, no significant results were observed for the time × microinjection interaction (F [8,520] = 1.34 ns) or the time × dose × microinjection interaction (F [24,520] = 1.37 ns).

### 3.3. Dopaminergic changes recorded in the *core* part of the nucleus accumbens following administration of ketamine (Figure 3)

In total 55 animals were used to investigate the ketamine effects on dopaminergic levels. Dopaminergic variations were found to be statistically dependent on the doses of ketamine and the postnatal microinjection. More specifically, statistical analysis showed that dopaminergic increases were dose-dependent and microinjection-dependent during the 90 min following saline and ketamine injections. Thus, the general ANOVA carried out for the 90 min post-injection displayed a significant dose-effect (NaCl 0.9%, ketamine 5 mg/kg, ketamine 10 mg/kg or ketamine 20 mg/kg) (F[3,47] = 9.64 p<0.00005), and a significant postnatal microinjection-effect (PBS/TTX) (F[1,47] = 16.34 p<0.0005), but no significant dose × microinjection interaction (F[3,47] = 1.44 n.s.).

**Figure 3.**
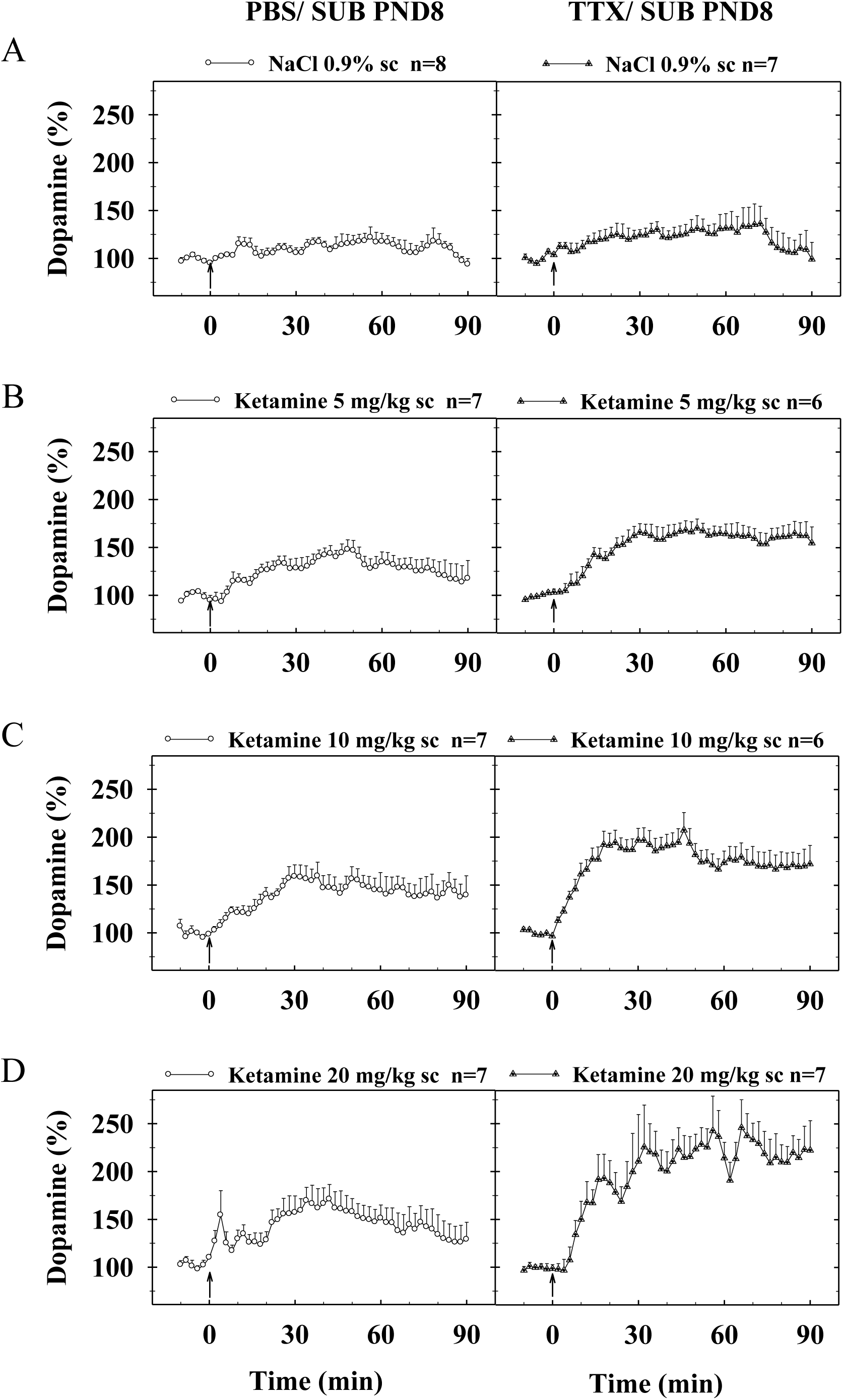
Ketamine-induced dopaminergic responses in the left *core* subregion of the nucleus accumbens of adult rats microinjected at postnatal day 8 (PND8) with PBS (left graphs) or TTX (right graphs) in the ventral subiculum (SUB). Animals received a subcutaneous (sc) injection (arrow) of NaCl 0.9% (A), ketamine 5 mg/kg (B), ketamine 10 mg/kg (C), or ketamine 20 mg/kg (D). Dopaminergic changes in the left *core* subregion of the nucleus accumbens were recorded every min using differential normal pulse voltammetry combined with computer-assisted numerical analysis of the voltammetric signal. Only mean values ± SEM corresponding to each two voltammograms are presented. Where no SEM is indicated, the size is less than the radius of the symbol. n is the number of animals in each group. Results are statistically analyzed using factorial ANOVA

The effects of ketamine on the time course of dopaminergic changes for the 90 min post-injection period as established by within-subjects analysis were as follows: a significant effect of time (F 44,2068] = 12.97 p < 0.00001), a significant time × dose interaction (F [132,2068] = 1.47 p < 0.001), a significant time × microinjection interaction (F [44,2068] = 2.56 p < 0.00001), and a significant time × dose × microinjection interaction (F [132,2068] = 1.54 p < 0.0005).

## 4. Discussion

The current study was devised within the framework of the animal modelling of the pathophysiology of schizophrenia. Its purpose was to determine the consequences of the transient postnatal functional blockade of the left ventral SUB for locomotor responses and DAergic responses measured in the *core* part of the Nacc following administration of the NMDA non-competitive antagonist, ketamine, in adult animals (rats). The *core* subregion of Nacc was targeted in the present study inasmuch as it has been found to be particularly involved in the pathophysiology of schizophrenia (McCollum et al., 2015) as well as in animal modelling studies performed after postnatal blockade of the prefrontal cortex (Meyer and Louilot, 2012; Tagliabue et al., 2017). In the present study, after postnatal inactivation of the SUB the results obtained following ketamine administration were enhanced locomotor reactivity, in particular to the dose 20 mg/kg sc, and increased and longer lasting DAergic variations.

Concerning locomotor activity, statistical analyses showed that responses during the 90 min following saline and ketamine injections were dose-dependent and postnatal microinjection-dependent. Interestingly, no differences were observed in locomotor activity in the period preceding the injection between the animals microinjected with PBS or TTX within the SUB, suggesting that the SUB neonatal inactivation has no significant impact on basal locomotor activity. As regards the animals microinjected with PBS in the SUB, the profile of the ketamine-induced responses, and in particular the marked response lasting about 50min for the 20 mg/kg sc ketamine dose, is consistent with the variations previously reported after neonatal injection of PBS in the prefrontal cortex (Usun et al., 2013; Pouvreau et al., 2016). In animals subjected to the neonatal TTX blockade of the SUB at PND8, ketamine-induced locomotor responses were more elevated and more durable for the two higher doses (10 mg/kg sc and 20 mg/kg sc), and in a significant way (compared to PBS animals) for the 20 mg/kg sc dose. As far as we know, the consequences of postnatal TTX SUB inactivation for ketamine-induced locomotor responses are being reported for the first time, making comparison with previous works difficult. However, it is interesting to note that the profile of locomotor hyperactivity observed in the present work is similar to the ketamine-induced locomotor responses observed after neonatal inactivation of the prefrontal cortex (Usun et al., 2013; Pouvreau et al., 2016), although not identical, inasmuch as maximum locomotor increases observed after neonatal SUB functional blockade with the 20 mg/kg sc ketamine dose appeared to be delayed in time. The reasons for that difference are yet to be determined.

Concerning the DAergic variations in the *core* part of the Nacc, statistical analyses revealed that increases during the 90 min following saline and ketamine injections were found to be dependent on the dose and on the neonatal microinjection in the SUB. Time-courses of these increases were also found to be dose-dependent and postnatal microinjection-dependent. To the best of our knowledge, it is the first time ketamine-induced DAergic changes have been reported in the *core* region of the Nacc *stricto sensu*. The DAergic increases observed in the animals microinjected with PBS in the SUB are compatible with the results of the meta-analysis published by Kokkinou et al. (2018) regarding the Nacc as a whole. In this respect, it is tempting to suggest that the high heterogeneity observed in the meta-analysis (Kokkinou et al., 2018) may have to do with opposite ketamine-induced DAergic variations in the shell and *core* subareas of the Nacc, inasmuch as such opposite variations have been observed in the two subregions after treatment with another non-specific NMDA antagonist, MK-801(Pouvreau et al., 2016; Tagliabue et al., 2017), and no differential regional analyses have been carried out in the other studies (Kokkinou et al., 2018). In animals microinjected with TTX the DA increases were higher and lasted longer than those observed in animals microinjected with PBS. Since it would appear that the ketamine-induced DAergic changes in the *core* part of the nucleus after TTX inactivation of the SUB, like the locomotor responses, have never previously been studied, direct comparison with other works is not possible.

Concerning the time-course of the two indexes, it seems that DAergic responses are delayed relative to the corresponding behavioural responses. This phenomenon is dependent on the particular doses of ketamine administered and was observed in animals microinjected in the SUB not only with PBS but also with TTX. Such a temporal mismatch between DAergic changes in the *core* part of the Nacc and locomotor responses has also been observed after administration of MK-801, another NMDA non-competitive antagonist, and neonatal TTX inactivation of the prefrontal cortex (Tagliabue et al., 2017). A similar temporal disconnection was also reported with administration (i.p.) of the well-known NMDA antagonist, phencyclidine (PCP), and DA detection in the Nacc with microdialysis (Adams and Moghaddam, 1998). With respect to the smallest dose of ketamine (5 mg/kg s.c.), the question arises as to whether the delayed DA variations may be due to the ketamine acting through its metabolites on AMPA receptors, insofar as it was recently reported that at low doses ketamine can have an antidepressant action through its metabolites and an NMDA receptor-independent mechanism (Zanos et al., 2016). Additional investigations failed to show an effect of the ketamine metabolites on stimulated DA release in the *core* part of the Nacc (Can et al., 2016), albeit these results were obtained in anesthetized animals and have yet to be confirmed in awake, freely moving animals, with other indexes of DA transmission. However, whereas given the available data in the literature the hypothesis of ketamine metabolite involvement in DA variations cannot be totally ruled out at this level, given the similarities between the temporal divergence between locomotor and DA variations observed with the different NMDA antagonists (ketamine, MK-801, PCP), which have different metabolites, it seems more likely that the ketamine effects we observed were linked to an action on NMDA receptors.

The temporal dissociation obtained in the present study between the time-courses of behavioural responses and DAergic responses suggests that the locomotor changes induced by ketamine correspond to an action on NMDA receptors that is at least partly independent of DA changes in the *core* part of the Nacc. However, it is important to emphasize that this does not mean ketamine effects on locomotor activity are not dependent on basal levels of dopamine in the Nacc. Indeed, it has been shown in rats (and the situation might be different in mice) that non-competitive NMDA antagonists are only able to restore locomotion when the dopaminergic system is moderately depleted or blocked, not when monoamine depletion is complete (Schmidt and Kretschmer, 1997). Moreover, as further suggested by these authors, if the anatomical target of glutamatergic antagonists to increase locomotor activity is downstream of the Nacc, such antagonists should be able to restore this behavioural response after the complete absence of striatal/accumbal DA, and yet this does not seem to be the case (Schmidt and Kretschmer, 1997). Insofar as stimulation of DA release in the *core* part of the Nacc by intra-accumbal local infusion of D-Amphetamine results in hyperlocomotor activity which is dependent on glutamatergic inputs in the Nacc (Rouillon et al., 2008), it is tempting to suggest that the impact of ketamine and, more generally, non-competitive NMDA antagonists on locomotor activity is dependent on a permissive role of DA, carried out by DA basal levels in the *core* part of Nacc. Other authors (Mele et al., 1998; De Leonibus et al., 2001; 2002) have proposed an alternative view, suggesting that locomotor hyperactivity induced by NMDA antagonists and locomotor activation induced by DA agonists (including D-Amphetamine) are mediated by different output pathways of the Nacc. It is true that two separate efferents pathways of the *core* part of the Nacc have been described, defined by medium spiny neurons possessing mainly D1 receptors (projecting to the substantia nigra/ventral tegmental area), and medium spiny neurons possessing mainly D2 receptors (projecting to the ventral pallidum) (see Humphries and Prescott, 2010). However, since all medium spiny neurons have been reported to possess NMDA receptors (Standaert et al., 1999), it is not easy to accommodate the proposal made by De Leonibus et al. (2001; 2002).

Whatever the case may be, it appears possible to suggest that the increased locomotor hyperactivity induced by ketamine observed in animals microinjected postnatally with TTX in the SUB is related to an increase in the sensitivity/density of NMDA receptors in the *core* part of Nacc. Direct glutamatergic projections from the SUB to the *core* part of Nacc have been described (French and Totterdell, 2002; Humphries and Prescott, 2010), offering the possibility that functional neonatal disconnection of this pathway could have an impact on glutamatergic transmission involving NMDA receptors in the *core* subregion. In this context it is also important to note that: 1) the SUB sends direct excitatory projections to the prefrontal cortex (Carr and Sesack, 1996; Witter, 2006); and 2) the functional inactivation of glutamatergic efferents of the prefrontal cortex in adult rats potentiates the increase in locomotor activity consecutive to local stimulation of DA transmission in the *core* part of Nacc (Rouillon et al., 2008). Thus, a second hypothesis would be that the behavioural results we obtained in the present study are also related to an indirect mechanism involving the prefrontal cortex, insofar as similar ketamine-induced behavioural responses were observed after postnatal prefrontal cortex inactivation (Usun et al., 2013; Pouvreau et al., 2016) and SUB inactivation (present study). However, convergent inputs of the prefrontal cortex and the SUB on the same neurons (medium spiny neurons) have been observed at the level of the *core* part of the Nacc (French and Totterdell, 2002), offering the possibility of coordination or interaction between the two input pathways in such a way that a dysfunctioning of one or other pathway would produce similar behavioural disruptions. In other words, it seems we cannot totally rule out the possibility that both a direct mechanism involving the SUB and an indirect mechanism involving the prefrontal cortex contributed to the present behavioural results obtained in animals microinjected with TTX in the SUB at PND8.

As regards DAergic responses in the *core* part of the Nacc, first of all in respect of the animals microinjected with PBS in the SUB at PND8, the most parsimonious hypothesis is to consider that the explanation for the DAergic variations induced by ketamine may be similar to that proposed for animals microinjected neonatally with PBS in the prefrontal cortex (Tagliabue et al., 2017). In short, in PBS animals, DA release induced by NMDA antagonists may result indirectly from an action on either NMDA receptors situated on intra-accumbal GABAergic interneurons, or GABAergic medium spiny neurons originating from the *core* subregion and projecting towards the ventral tegmental area (for a detailed discussion see Tagliabue et al., 2017). Concerning the animals microinjected postnatally with TTX, and assuming that enhancement of locomotor hyperactivity induced by ketamine in these animals is related to an increased sensitivity/density of NMDA receptors situated in the *core* part of the Nacc, it is tempting to suggest that bigger DA increases in the *core* part of the Nacc in animals microinjected postnatally with TTX are also related to similar changes involving increased sensitivity/density of NMDA receptors. More precisely, this is a suggestion consistent with the existence of synaptic terminals derived from direct glutamatergic projection of the SUB to neurons in the *core* part of Nacc (French and Totterdell, 2002). It is also important to note here that, in contrast to most forebrain regions, there have been no reports of direct projections from the SUB to the ventral tegmental area, the area of origin of DA neurons (Geisler and Zahm, 2005), which suggests that the neonatal SUB inactivation would not affect NMDA receptors in the ventral tegmental area. It is worth mentioning that it has been reported that stimulation of the SUB in *adults* rats increased the number of spontaneously active DA neurons in the ventral tegmental area, and that this increase was completely eliminated after glutamate receptor blockade in the Nacc and independent of the prefrontal cortex (Floresco et al., 2001). More recently, however, this control has been shown to concern only the *shell* subregion of the Nacc, not the *core* subregion (Peleig-Rabstein and Feldon; 2006). Furthermore, although this phenomenon has been described in normal adults, the situation after neonatal inactivation of the SUB may be different. Interestingly, the neonatal lesion of the ventral hippocampus has been found to lead to alterations of neuronal arborization and spine density in the prefrontal cortex (Flores et al., 2005), as well as functional disruption of this forebrain structure (Macedo et al., 2012). In other respects, the prefrontal cortex sends marked projections to the *core* part of the Nacc (French and Totterdell, 2002; Humphries and Prescott, 2010). Moreover, a convergence of terminals derived from the prefrontal cortex and the SUB was observed on the distal dendrites of identified medium-sized, densely spiny neurons (French and Totterdell, 2002). Thus, it is possible to propose, somewhat tentatively, that after postnatal TTX inactivation of the SUB, the prefrontal cortex was also involved, albeit indirectly, in DA increases in the *core* part of the Nacc following ketamine administration.

Finally, it is important to note that the hypothesis proposed to explain the behavioural and DAergic results we obtained, and according to which the sensitivity/density of NMDA receptors changes after SUB TTX postnatal blockade, is compatible with data showing that NMDA receptors are subjected to conformational changes during the 3 postnatal weeks (see Stroebel et al., 2018), and that these conformational changes are dependent on neuronal activity, with neuronal blockade by TTX increasing the pool of NR1/NR2B NMDA postsynaptic receptors (Perez-Otana and Ehlers, 2005). If we now consider human post-mortem data obtained from patients with schizophrenia, no differences were reported in NMDA expression or binding in the Nacc (Noga et al., 1997; Meador-Woodruff et al., 2001; Aparicio-Legarza et al., 1998). However, in the afore-mentioned studies no *core* and *shell* subdivisions of the Nacc were reported, which deserves further research.

## 5. Conclusion

In conclusion, what is interesting in the present study is that the behavioural changes observed after ketamine administration precede the dopaminergic changes in the *core* part of the Nacc. The question that may be asked here is: Does this process occur with the development of schizophrenia? In other words, if we accept that NMDA transmission is disrupted in schizophrenia, as the results obtained in humans with NMDA antagonists suggest, it is tempting to propose that behavioural disturbances characteristic of the pathophysiology of schizophrenia occur before perturbations of the DAergic transmission. In other respects, it is also interesting to note that many years ago it was suggested that there is a positive functional interdependence between DA neurons reaching, on the one hand, the Nacc and, on the other hand, the dorsal striatum (Louilot et al., 1987b). Therefore, it is tempting to suggest that a primary impairment of DA transmission in the *core* part of the Nacc may ultimately contribute to the elevated presynaptic DAergic disturbance recently observed in the dorsal striatum (Weinstein et al., 2017). Thus, the involvement of NMDA receptors’ functional disruption in the *core* part of the Nacc in the pathophysiology of schizophrenia warrants further investigation.

## Acknowledgements

The authors are grateful to Tiphaine Pouvreau for her helpful assistance. The authors wish also to thank Elora Kereselidze for her help in preparing the final version of the manuscript.

## References

Adams, B., Moghaddam, B., 1998. Corticolimbic dopamine neurotransmission is temporally dissociated from the cognitive and locomotor effects of phencyclidine. J. Neurosci. 18, 5545–5554.

Aparicio-Legarza, M.I., Davis, B., Hutson, P.H, Reynolds, G.P., 1998. Increased density of glutamate/N-methyl-D-aspartate receptors in putamen from schizophrenic patients. Neurosci. Lett. 241, 143–146.

Arnold, S.E., Rioux, L., 2001. Challenges, status, and opportunities for studying developmental neuropathology in adult schizophrenia. Schizophr. Bull. 27, 395–416.

Cao, H., Dixson, L., Meyer-Lindenberg, A., Tost, H., 2016, Functional connectivity measures as schizophrenia intermediate phenotypes: advances, limitations, and future directions. Curr Opin Neurobiol. 36, 7–14.

Can, A., Zanos, P., Moaddel, R., Kang, H.J., Dossou, K.S., Wainer, I.W., Cheer, J.F., Frost, D.O., Huang, X.P., Gould, T.D., 2016. Effects of Ketamine and Ketamine Metabolites on Evoked Striatal Dopamine Release, Dopamine Receptors, and Monoamine Transporters. J. Pharmacol. Exp. Ther. 359, 159–170.

Carr, D.B., Sesack, S.R., 1996. Hippocampal afferents to the rat prefrontal cortex: synaptic targets and relation to dopamine terminals. J. Comp. Neurol. 369, 1–15.

Coyle JT, Basu A, Benneyworth M, Balu D, Konopaske G., 2012, Glutamatergic synaptic dysregulation in schizophrenia: therapeutic implications. Handb Exp Pharmacol. 213, 267–295.

De Leonibus, E., Mele, A., Oliverio, A., Pert, A., 2001. Locomotor activity induced by the non-competitive N-methyl-D-aspartate antagonist, MK-801: role of nucleus accumbens efferent pathways. Neuroscience 104, 105–116.

De Leonibus, E., Mele, A., Oliverio, A., Pert, A., 2002. Distinct pattern of c-fos mRNA expression after systemic and intra-accumbens amphetamine and MK-801. Neuroscience 115, 67–78.

Field, K.J, White, W.J., Lang, C.M., 1993. Anaesthetic effects of chloral hydrate, pentobarbitone and urethane in adult male rats. Lab. Anim. 27, 258–269.

Floresco, S.B., Todd, C.L., Grace, A.A., 2001. Glutamatergic afferents from the hippocampus to the nucleus accumbens regulate activity of ventral tegmental area dopamine neurons. J Neurosci. 21, 4915–4922.

Flores, G., Alquicer, G., Silva-Gómez, A.B., Zaldivar, G., Stewart, J., Quirion, R., Srivastava, L.K., 2005 Alterations in dendritic morphology of prefrontal cortical and nucleus accumbens neurons in post-pubertal rats after neonatal excitotoxic lesions of the ventral hippocampus. Neuroscience 133, 463–470.

Fornito, A., Bullmore, E.T., 2015. Reconciling abnormalities of brain network structure and function in schizophrenia. Curr. Opin. Neurobiol. 30, 44–50.

French, S.J, Totterdell, S., 2002. Hippocampal and prefrontal cortical inputs monosynaptically converge with individual projection neurons of the nucleus accumbens. J. Comp. Neurol. 446, 151–165.

Friston, K.J., Frith, C.D., 1995. Schizophrenia: a disconnection syndrome? Clin. Neurosci. 3, 89–97.

Geisler, S., Zahm, D.S., 2005 Afferents of the ventral tegmental area in the rat-anatomical substratum for integrative functions. J. Comp. Neurol. 490, 270–294.

Gonzalez-Mora, J.L., Guadalupe, T., Fumero, B., Mas, M., 1991. Mathematical resolution of mixed in vivo voltammetry signals. Models, equipment, assessment by simultaneous microdialysis sampling. J. Neurosci. Methods 39, 231–244.

Heimer, L., Alheid, G.F., de Olmos, J.S., Groenewegen, H.J., Haber, S.N., Harlan, R.E., Zahm, D.S., 1997. The accumbens: beyond the core-shell dichotomy. J. Neuropsychiatry Clin. Neurosci. 9, 354–381.

Humphries, M.D., Prescott, T.J., 2010. The ventral basal ganglia, a selection mechanism at the crossroads of space, strategy, and reward. Prog. Neurobiol. 90, 385–417.

Inta, D., Meyer-Lindenberg, A., Gass, P., 2011. Alterations in postnatal neurogenesis and dopamine dysregulation in schizophrenia: a hypothesis. Schizophr Bull. 37, 674–680.

Kokkinou, M., Ashok, A.H., Howes, O.D., 2018. The effects of ketamine on dopaminergic function: meta-analysis and review of the implications for neuropsychiatric disorders. Mol. Psychiatry 23, 59–69.

Lahti, A.C., Weiler, M.A., Michaelidis, T., Parwani, A., Tamminga, C.A., 2001. Effects of ketamine in normal and schizophrenic volunteers. Neuropsychopharmacology 25, 455–467.

Louilot, A., Serrano, A., D’Angio, M., 1987a. A novel carbon fiber implantation assembly for cerebral voltammetric measurements in freely moving rats. Physiol. Behav. 41, 227–231.

Louilot, A., Taghzouti, K., Deminiere, J.M., Simon, H., Le Moal, M., 1987b. Dopamine and behavior: functional and theoretical considerations in: Sandler, M., Feuerstein, C., Scatton, B.(Eds), Neurotransmitters interactions. Raven Press, New-York, pp. 193–204.

Macedo, C.E., Angst, M.J., Gobaille, S., Schleef, C., Guignard, B., Guiberteau, T., Louilot, A., Sandner, G., 2012. Prefrontal dopamine release and sensory-specific satiety altered in rats with neonatal ventral hippocampal lesions. Behav. Brain Res. 231, 97–104.

McCollum, L.A., Walker, C.K., Roche, J.K., Roberts, R.C., 2015. Elevated Excitatory Input to the Nucleus Accumbens in Schizophrenia: A Postmortem Ultrastructural Study. Schizophr Bull. 41, 1123–1132.

Meador-Woodruff, J.H., Hogg, A.J. Jr., Smith, R.E., 2001. Striatal ionotropic glutamate receptor expression in schizophrenia, bipolar disorder, and major depressive disorder. Brain Res. Bull. 55, 631–640.

Mele, A., Thomas, D.N., Pert, A., 1998. Different neural mechanisms underlie dizocilpine maleate- and dopamine agonist-induced locomotor activity. Neuroscience 82, 43–58.

Meyer, F., Peterschmitt, Y., Louilot, A., 2009. Postnatal functional inactivation of the entorhinal cortex or ventral subiculum has different consequences for latent inhibition-related striatal dopaminergic responses in adult rats. Eur. J. Neurosci. 29, 2035–2048.

Meyer, F., Louilot, A., 2011. Latent inhibition-related dopaminergic responses in the nucleus accumbens are disrupted following neonatal transient inactivation of the ventral subiculum. Neuropsychopharmacology 36, 1421–1432.

Meyer, F., Louilot, A., 2012. Early prefrontal functional blockade in rats results in schizophrenia-related anomalies in behavior and dopamine. Neuropsychopharmacology 37, 2233–2243.

Meyer, F., Louilot, A., 2014. Consequences at adulthood of transient inactivation of the parahippocampal and prefrontal regions during early development: new insights from a disconnection animal model for schizophrenia. Front. Behav. Neurosci. 8, 118.

Moczydlowski, E.G., 2013. The molecular mystique of tetrodotoxin. Toxicon. 1, 165–183.

Moghaddam, B., Adams, B.W., 1998. Reversal of phencyclidine effects by a group II metabotropic glutamate receptor agonist in rats. Science 281, 1349–1352.

Noga, J.T., Hyde, T.M., Herman, M.M., Spurney, C.F., Bigelow, L.B., Weinberger, D.R., Kleinman, J.E., 1997. Glutamate receptors in the postmortem striatum of schizophrenic, suicide, and control brains. Synapse 27, 168–176.

Owen, M.J, Sawa, A., Mortensen, P.B., 2016. Schizophrenia. Lancet. 388, 86–97.

Paxinos, G., Watson, C., 2009. The rat brain in stereotaxic coordinates, compact sixth edition. Academic Press, London.

Peleg-Raibstein, D., Feldon, J., 2006. Effects of dorsal and ventral hippocampal NMDA stimulation on nucleus accumbens core and shell dopamine release. Neuropharmacology 51, 947–957.

Pérez-Otaño, I., Ehlers, M.D., 2005. Homeostatic plasticity and NMDA receptor trafficking. Trends Neurosci. 28, 229–238.

Piper, M., Beneyto, M., Burne, T.H., Eyles, D.W., Lewis, D.A., McGrath, J.J., 2012. The neurodevelopmental hypothesis of schizophrenia: convergent clues from epidemiology and neuropathology. Psychiatr. Clin. North Am. 35, 571–584.

Pouvreau, T., Tagliabue, E., Usun, Y., Eybrard, S., Meyer, F., Louilot, A., 2016. Neonatal Prefrontal Inactivation Results in Reversed Dopaminergic Responses in the Shell Subregion of the Nucleus Accumbens to NMDA Antagonists. ACS. Chem. Neurosci. 7, 964–971.

Rosenthal, R., Rosnow, R.L., Rubin, D.B., 2000. Contrasts and effect sizes in behavioral research: a correlational approach. Cambridge University Press, New York.

Rouillon, C., Abraini, J.H., David, H.N., 2008. Prefrontal cortex and basolateral amygdala modulation of dopamine-mediated locomotion in the nucleus accumbens core. Exp. Neurol. 212, 213–217.

Saha, S., Chant, D., Welham, J., McGrath, J., 2005. A systematic review of the prevalence of schizophrenia. PLoS Med. 2, e141.

Schmidt, W.J., Kretschmer, B.D., 1997. Behavioural pharmacology of glutamate receptors in the basal ganglia. Neurosci. Biobehav. Rev. 21, 381–392.

Schobel, S.A., Chaudhury, N.H., Khan, U.A., Paniagua, B., Styner, M.A., Asllani, I., Inbar, B.P., Corcoran, C.M., Lieberman, J.A., Moore, H., Small, S.A., 2013. Imaging patients with psychosis and a mouse model establishes a spreading pattern of hippocampal dysfunction and implicates glutamate as a driver. Neuron 78, 81–93.

Standaert, D.G., Friberg, I.K., Landwehrmeyer, G.B., Young, A.B., Penney, J.B. Jr., 1999. Expression of NMDA glutamate receptor subunit mRNAs in neurochemically identified projection and interneurons in the striatum of the rat. Mol. Brain Res. 64, 11–23.

Stroebel, D., Casado, M., Paoletti, P., 2018. Triheteromeric NMDA receptors: from structure to synaptic physiology. Curr. Opin. Physiol. 2, 1–12.

Tagliabue, E., Pouvreau, T., Eybrard, S., Meyer, F., Louilot, A., 2017. Dopaminergic responses in the core part of the nucleus accumbens to subcutaneous MK801 administration are increased following postnatal transient blockade of the prefrontal cortex. Behav. Brain. Res. 335, 191–198.

Usun, Y., Eybrard, S., Meyer, F., Louilot, A., 2013. Ketamine increases striatal dopamine release and hyperlocomotion in adult rats after postnatal functional blockade of the prefrontal cortex. Behav. Brain Res. 256, 229–237.

Weinberger, D.R., 1995. From neuropathology to neurodevelopment. Lancet. 346, 552–557.

Weinstein, J.J., Muhammad, O.C., Slifstein, M., Kegeles, L.S., Moore, H., Abi-Dargham, A., 2017. Pathway-specific dopamine abnormalities in schizophrenia. Biol. Psychiatry 81, 31–42.

Winship, I.R., Dursun, S.M., Baker, G.B., Balista, P.A., Kandratavicius, L., Maia-de Oliveira, J.P., Hallak, J., Howland, J.G., 2019. An overview of animal models related to schizophrenia. Can. J. Psychiatry 64, 5–17.

Witter, M.P., 2006 Connections of the subiculum of the rat: Topography in relation to columnar and laminar organization. Behav. Brain Res. 174, 251–264.

Zanos, P., Moaddel, R., Morris, P.J., Georgiou, P., Fischell, J., Elmer, G.I., Alkondon, M., Yuan, P., Pribut, H.J., Singh, N.S., Dossou, K.S., Fang, Y., Huang, X.P., Mayo, C.L., Wainer, I.W., Albuquerque, E.X., Thompson, S.M., Thomas, C.J., Zarate, C.A. Jr, Gould, T.D., 2016. NMDAR inhibition-independent antidepressant actions of ketamine metabolites. Nature 533, 481–486.

